# Contextual protein and antibody encodings from equivariant graph transformers

**DOI:** 10.1101/2023.07.15.549154

**Authors:** Sai Pooja Mahajan, Jeffrey A. Ruffolo, Jeffrey J. Gray

## Abstract

The optimal residue identity at each position in a protein is determined by its structural, evolutionary, and functional context. We seek to learn the representation space of the optimal amino-acid residue in different structural contexts in proteins. Inspired by masked language modeling (MLM), our training aims to transduce learning of amino-acid labels from non-masked residues to masked residues in their structural environments and from general (e.g., a residue in a protein) to specific contexts (e.g., a residue at the interface of a protein or antibody complex). Our results on native sequence recovery and forward folding with AlphaFold2 suggest that the amino acid label for a protein residue may be determined from its structural context alone (i.e., without knowledge of the sequence labels of surrounding residues). We further find that the sequence space sampled from our masked models recapitulate the evolutionary sequence neighborhood of the wildtype sequence. Remarkably, the sequences conditioned on highly plastic structures recapitulate the conformational flexibility encoded in the structures. Furthermore, maximum-likelihood interfaces designed with masked models recapitulate wildtype binding energies for a wide range of protein interfaces and binding strengths. We also propose and compare fine-tuning strategies to train models for designing CDR loops of antibodies in the structural context of the antibody-antigen interface by leveraging structural databases for proteins, antibodies (synthetic and experimental) and protein-protein complexes. We show that pretraining on more general contexts improves native sequence recovery for antibody CDR loops, especially for the hypervariable CDR H3, while fine-tuning helps to preserve patterns observed in special contexts.

## Introduction

### Motivation

The physical forces that drive residue-residue interactions in proteins, at protein-protein interfaces and at antibody-antigen interfaces are the same. However, it is the context of the residue, structural, evolutionary and functional, that ultimately determines the optimal amino acid at a given position in a protein. Some successful protein DL models including AlphaFold2 and ProGen2 have been found to perform more weakly in certain important contexts such as antibodies.^1, 2^ Yet some important contexts, like TCR modeling,^3^ suffer from a lack of data. An open question is when it is best to use general protein models, and when it is best to use models for specific contexts, or perhaps how to combine general and specific data to learn effectively for different contexts. Here, we apply masked label prediction, an extension of the ideas introduced Shi et al.^4^ with **M**asked **L**anguage **M**odeling^5^ (MLM) for graph structured data, to learn encodings for the optimal amino-acid residue in different contexts.

MLM was first introduced with the Bidirectional Encodings from Transformers^5^ (BERT) model to pretrain deep bidirectional representations from unlabeled text by jointly conditioning on both left and right context in text-based data. In MLM, ∼15% of the words in the input are randomly masked (identity is hidden by replacing it with a token), and the entire text sequence is run through the attention-based transformer encoder and *then the masked words are predicted based on the context provided by the non-masked words in the sequence.* Since its introduction in 2019, MLM has been applied to pretrain models for a variety of texts and to fine-tune on a wide range of tasks^6^.

Shi *et al.* proposed UniMP^4^, a BERT-inspired masked label prediction model for graph structured data that *simulates the procedure of transducing label information from labeled to unlabeled examples in the graph*. During training, a certain percentage of labels are masked, and UniMP propagates both feature and label information, and the model learns to predict the masked labels based on the information from the non-masked labels.

Here, we set out to apply the conceptual framework of MLM^5^ and UniMP^4^ to *transduce learning of amino-acid residue labels* from **a)** non-masked amino-acid residues in their structural and sequence environments to masked amino-acid residues in their structural and partially (< 100%) or fully masked (100% masking) sequence environments (or structure-only environments) and **b)** more-general contexts (a residue in a protein) to more-specific contexts (a residue at the interface of a heteromeric protein or at the antibody-antigen interface) (**Figure 1A**), similar to fine-tuned BERT models on domain-specific data.

There are several questions we can probe. For aim (a), we investigate the ability of different models to predict the identity of an amino-acid residue (native sequence recovery), in the absence of any sequence information (100% masking i.e., from its structural context alone) for a model trained to transduce labels in a partially masked (15-25%) context (**Figure 1A, B**).

For aim (b), we compare models trained by masking amino-acid residues in different contexts for native sequence recovery with different pretraining and fine-tuning strategies (**Figure 1C**). To understand the motivation behind this approach, consider the task of designing the complementarity determining regions (CDRs) of an antibody that recognize and bind an antigen of interest. This task can be framed as predicting the identity of amino-acid residues of the CDR loops of the antibody in the context of the antigen at the antibody-antigen interface. This context is specialized since the amino-acid residues in antibodies exhibit distinct conservation patterns but also must vary sufficiently to confer shape and chemical complementarity to the epitope on the antigen. We hypothesized that such a task could benefit from pretraining on the more general contexts of proteins, and protein-protein interfaces. In other words, the three-dimensional association patterns formed by the 20 amino acids in these more general contexts must bear some resemblance to those at the antibody-antigen interface. Furthermore, for this specific data-sparse task, pretraining on more general contexts allows us to leverage large corpuses of structural data for proteins, protein-protein interfaces and antibodies instead of just relying on the smaller datasets for antibody-antigen interfaces.

Overall, we seek to investigate whether contextual encodings derived through pretraining and fine-tuning (in a manner similar to BERT models in natural language processing) are advantageous for representing different structural and evolutionary contexts in proteins, particularly the complementarity determining regions in the CDR loops of antibodies. We further investigate the sequence and structural space sampled by our masked models in different contexts for various desirable properties such as folding into target structures, recapitulating wildtype binding energies at interfaces and for retaining conserved sequence patterns in antibody designs at the antibody-antigen interface while also learning the CDR mutations driven by the antigen.

### Related Work

Jing et al.^7^ focused on learning sequence and structural representations (dense, all-atom representation) simultaneously for a range of tasks on the ATOM3D dataset (such as predicting binding, non-binding residues, small molecule binding etc.). Other important studies that apply neural networks to encode sequence and three-dimensional structures for representation learning include GearNet^8^ and ESM-GearNet^9^ that focused primarily on EC and GO-prediction tasks and MIF-ST^10^ that focused on zero-shot stability prediction. We aim to learn coarse representations (**Figure 1B**) for zero-shot sequence generation from masked representations in different physical and biological contexts of interest.

When the model is provided structural context alone (100% masking at inference), our model predicts pseudolikelihood of amino-acid labels at each position from structure information alone similar to “inverse folding” models such as the Structured Graph transformer^11^, GVP^12^, ESM-IF1^13^ and ProteinMPNN^14^. However, our model (**Figure 1B**) is distinct from these models that comprise a structure-only encoder and rely on a sequence decoder (autoregressive or otherwise) to generate the sequence given the structure both during training and inference. Our model, on the other hand, learns masked sequence labels in the context of the sequence and structure of the unmasked nodes similar to MIF-ST^10^ and PiFold^15^. We further focus questions on whether such contextual encodings derived through pretraining and fine-tuning (in a manner similar to BERT models) learn representations for specialized contexts such as the antibody-antigen interface. While others have investigated the effect of pretraining and fine-tuning on protein families for protein language models (ESM-v1^16^, ProGen2^1^), we are interested in asking this question for sequence-structure encoders.

## Results and discussion

### Masked label prediction given 3D structural and sequence context

Our model is inspired by the BERT model’s architecture that is a multi-layer bidirectional Transformer encoder (**Figure 1B**). Since we represent protein structure as a graph, our encoder consists of four E(n)Transformer blocks based on the Equivariant Graph Neural Network^17^ (EGNN) and modified to include transformer-like layers for attention-based aggregation over neighboring nodes, feedforward layers and multi-head attention (see Methods). Each residue is represented by three backbone atoms (N, C_A_, C) and one sidechain atom (C_B_). Node features consist of twenty one-hot encoded sequence labels (amino acid residues) and four one-hot encoded atom labels (N, C_A_, C, C_B_). All amino-acid residues, including glycine, are appropriately represented by such a representation (C_B_ is encoded as a 0 for glycine). For masked residues, C_B_ is removed (one-hot atom encoding set to zero, same as glycine) and its coordinates are replaced by zeros. The relative positions in space and residue positions in the sequence are encoded with Roformer^24^ and sinusoidal embeddings^25^ respectively. When multiple chains are present, positional indices are offset by 100 residues. Each atom is connected to its 48 nearest neighbor atoms. We use a hidden dimension (D) of 256 split over 8 attention heads.

**Figure 1.**
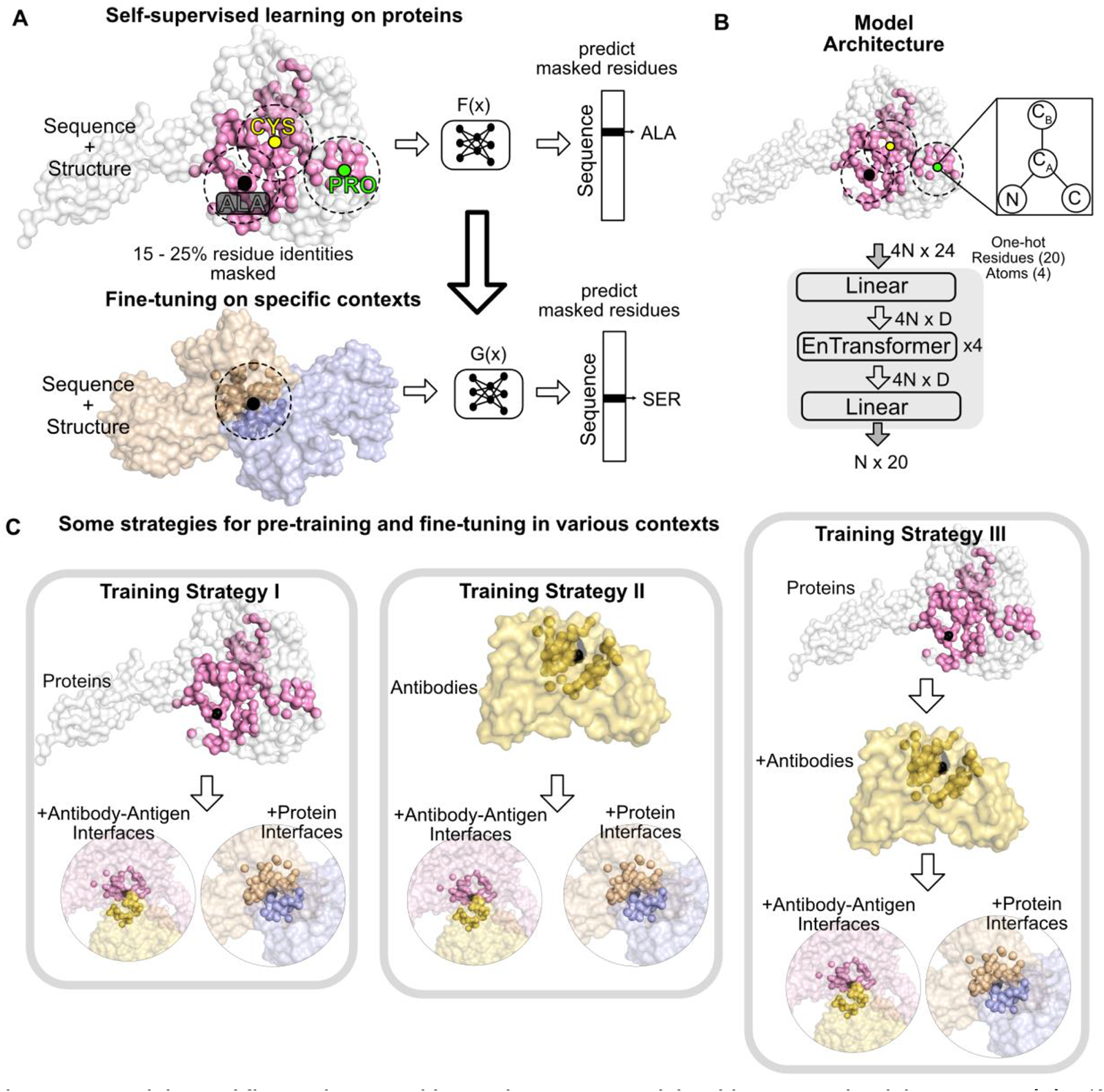
Pretraining and fine-tuning on residue environments, model architecture and training strategy. **(A)** Self-supervised learning to transduce sequence labels for masked residues (e.g., ALA) from those for unmasked residues (CYS, PRO) by context matching on proteins. The encoding thus learned can be fine-tuned to specialized contexts such as protein-protein interfaces (shown) or CDR loops of antibodies (not shown) at the antibody-antigen interface. **(B)** Model architecture for encoding the structural and sequence context of amino-acid residues. The model utilizes an equivariant GNN (EGNN^17^) with attention to aggregate node information from neighboring nodes (atoms). The model with hidden dimension *D*, inputs one-hot sequence and atom information (N, C_A_, C and C_B_) and the connectivity between the nodes in 3D space for a protein with *N* amino acid residues to predict the sequence label at masked positions. **(C)** We propose three training strategies for pretraining on general contexts and fine-tuning on specialized contexts. In training strategy I (left), we train the model on all single chains in the PDB50^18, 19^ dataset followed by fine-tuning on residues at protein-protein interfaces (MaSIF^20^) followed by fine-tuning on residues at the antibody-antigen interfaces. In training strategy II (center), we train the model on the SAbDAb^21^ and AF2 antibody datasets^22^ followed by fine-tuning simultaneously on the antibody-antigen interfaces and the protein-protein interfaces. In training strategy III (right), we train the model on all single chains in the PDB50^18,19^ dataset followed by fine-tuning on a dataset of antibody structures from SAbDAb^21^ and AlphaFold2^23^ (AF2) followed by finetuning simultaneously on the antibody-antigen interfaces and the protein-protein interfaces.

#### Strategy for pretraining and fine-tuning context-aware models

To train a model that is both context-aware (sequence and structure) and leverages information from the general context, we follow a hierarchical training strategy (**Figure 1C**). Furthermore, to prevent the fine-tuned model from catastrophic forgetting, a phenomenon in which a fine-tuned neural network forgets information learnt from pre-training, we subsample data points from the pretraining datasets during fine-tuning.

We investigate three training strategies for pretraining on general contexts and fine-tuning on specialized contexts (**Figure 1C**). In “training strategy I”, we train the model on all single chains in the PDB50^18, 19^ dataset followed by fine-tuning on residues at protein-protein interfaces (MaSIF^20^ structurally diverse dataset or DIPS heterodimers-only dataset) followed by fine-tuning on residues at the antibody-antigen interfaces. In training strategy II, we train the model on all single chains in the PDB50 dataset followed by fine-tuning on a dataset of antibody structures from SAbDAb^21^ and AlphaFold2^23^ (AF2) followed by fine-tuning simultaneously on the antibody-antigen interfaces and the protein-protein interfaces. In training strategy III, we omit pretraining on the PDB50 set and train the model on the SAbDAb and AF2 antibody datasets followed by fine-tuning simultaneously on the antibody-antigen interfaces and the protein-protein interfaces (details in Methods).

#### Model generalizes sequence label prediction from partially masked context in training to fully masked context at inference

In specialized biological contexts, such as those for residues on the antibody paratope, the structural and sequence context must be determined from the sequence and structural context of the binding interface. Thus, a model that learns partially masked representations is relevant in such a context.

To this end, we determined the ability of a model trained with the masked label objective (Figure 1A, 1B) to transduce learning from lower mask rates (15%-25%) at training to higher mask rates at inference. We test this for ProtEnT, a model trained on proteins and the basis for pretraining strategies I and III. For this model, we calculated the sequence recovery on the TS50 test set as a function of different masking rates and temperatures (**Figure 2A** and **SI Figure 1**). We find that while the sequence recovery drops as higher percentage of residues are masked, even at 100% masking, the model can recover 37.4% residues with structure information alone and without additional decoding (one-shot sequence prediction). In later sections, we probe sequences generated at 100% masking for folding and overlap with natural sequences.

For intermediate masking rates (40 – 60%) that are relevant for predicting sequence identity of the residues at the interface in the context of a binding partner, the model can recover up to 44% of the native sequence (temperatures 0.01 – 0.1). The relatively high native sequence recovery in this work and another recent study^15^ suggests that it may be possible to predict amino acid identity given full structural and partial sequence context in one-shot, i.e., without a sequence decoder.

#### Pretraining and contextual fine-tuning improves native sequence recovery

To determine whether pretraining on protein and/or protein complexes improves sequence recovery on specialized and data-limited contexts such as the antibody-antigen interface (training strategy I; Figure 1C), we obtained the sequence recovery for models trained with different training schemes (**Table 1**). The model names for baseline, pretrained and fine-tuned models are listed in Table 1. For example, ProtEnT is trained on proteins only, PPIEnT is trained on protein-protein complexes only, ProtAbAgEnT is pretrained on proteins followed by fine-tuning on antibody-antigen complexes and so on.

Comparing the performance of ProtEnT, PPIEnT and ProtPPIEnT models on the PPI300 test set, we find that fine-tuning on protein interfaces (ProtPPIEnT) shows only a small improvement in sequence recovery over the ProtEnT model. This suggests that the residue environments at protein interfaces resemble those at the core of proteins and can be recovered accurately by models trained on proteins alone. This observation is supported by other studies by Chu et al.^26^ and Dauparas et al.^14^ who reported high sequence recoveries for protein complexes for models trained on protein structures.

For sequence recovery on the AbAg70 test set, we find that pretraining leads to significant improvement. Surprisingly, pretraining on proteins and fine-tuning on antibody-antigen complexes (ProtAbAgEnT) leads to the most pronounced improvement in sequence recovery over the AbAgEnT (33.3% to 46.9%) and the ProtEnT (38.4% to 46.9%) models. The best performing model is trained with data from all three contexts (ProtPPIAbAgEnT) though the improvement is not significant over the ProtAbAgEnT. While the PPIEnT model trained only on interface residues in protein complexes (excluding antibody complexes) *does not* benefit the sequence recovery at antibody-antigen interfaces (see more details in following sections), training on both protein complexes and antibody complexes (PPIAbAgEnT) boosts sequence recovery over the model trained on antibody complexes alone (AbAgEnT).

**Table 1.** Sequence recovery with models from different training strategies models on protein, protein-protein and antibody-antigen test sets. Sequency recovery is calculated at 100% masking protein dataset. For protein-protein and antibody-antigen datasets, we mask and predict *interface* residues on *one* binding partner (masked) in the context of the other partner (not masked).

|  | Training |  |  |  | Sequence Recovery |  |  |
| --- | --- | --- | --- | --- | --- | --- | --- |
|  |  | Protein | PPI | AbAg | ProteinsTS50 | PPI300 | AbAg70 |
| baseline | ProtEnT | X |  |  | **37.6±6.0** | 40.7±7.8 | 38.4±6.5 |
|  | PPIEnT |  | X |  | 26.8±4.0 | 32.0±7.0 | 32.5±7.0 |
|  | AbAgEnT |  |  | X | 12.2±3.0 | 15.0±4.6 | 33.3±11.0 |
|  | PPIAbAgEnT |  | X | X | 17.4±4.0 | 25.8±6.3 | 37.0±9.0 |
| with pretraining | ProtPPIEnT | X | X |  | 35.3±5.0 | **41.9±8.0** | 37.3±6.5 |
|  | ProtPPIAbAgEnT | X | X | X | 34.5±5.0 | 41.1±8.0 | **47.1±8.0** |
|  | ProtAbAgEnT | X |  | X | 32.5±5.0 | 39.1±7.2 | **46.9±9.0** |

**Figure 2.**
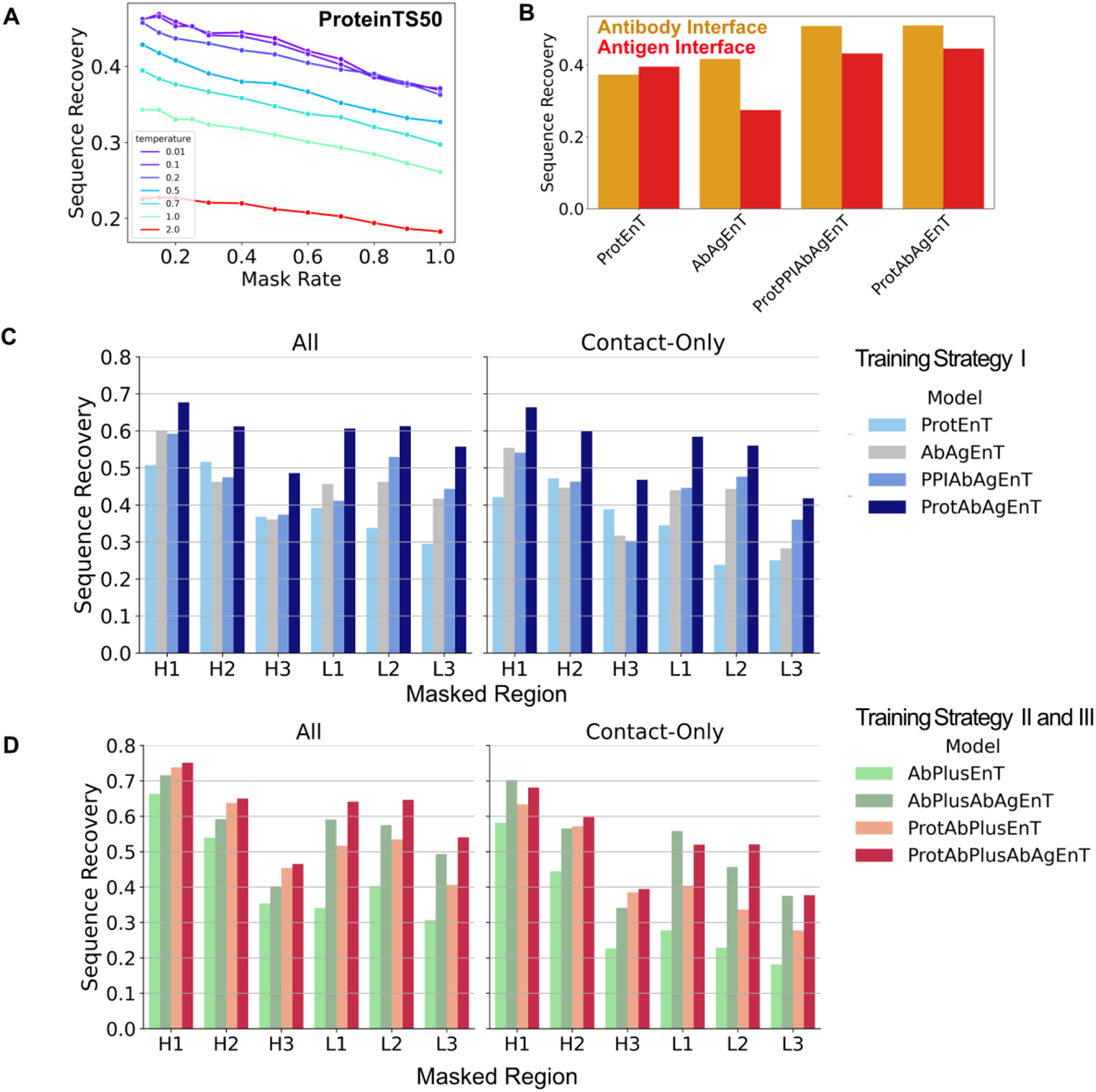
Comparison of sequence recovery on test sets with different masking and sampling schemes. **(A)** Sequence recovery as a function of percentage masking and sampling temperature during inference. Sequence recovery for the ProtEnT model on the ProteinTS50 test set for temperatures ranging from 0.01 to 2.0 is plotted against inference masking percentage ranging from 10% to 100%. The model was trained with a 15% masking rate. **(B)** Sequence recovery for the AbAg70 test set for four representative models from training strategy I. The sequence recovery is broken down by the recovery on the masked antibody or antigen interface. The reported recovery is the average recovery on the test set over 100 sequences sampled per each target. **(C)** and **(D)** Sequence recovery on the AbAg70 test set for either a masked CDR loop (left subplot) or only masked contact residues on the CDR loop (right subplot). For left subplots, we report median sampled recovery over 100 sampled sequences for each target. For the right subplot, since the number of contact residues can vary significantly per loop per target, we report an average recovery over the full dataset (i.e. (total matches with native sequence) / (total contact residues sampled per loop)). In (C) we show plots for training strategy I and in (D) we show the same plots for training strategies II (greens) and III (reds).

#### Fine-tuned models retain accuracy on proteins and protein interfaces

To test whether the successive training and subsampling strategy proposed in this work helps the fine-tuned models retain sequence recovery on generalized datasets, we calculated the sequence recovery for fine-tuned models on datasets they were pretrained on.

The fine-tuned ProtPPIEnT model achieves a sequence recovery of 41.9% on the PPI300 test set while retaining sequence recovery of 35.3% on the ProteinsTS50 test set. Similarly, the ProtPPIAbAgEnT retains high sequence recoveries on the ProteinsTS50 (34.5%) and PPI300 (41.1%) datasets. Thus, the fine-tuned models are context-aware and still general.

#### Fine-tuning on antibody-antigen interfaces improves sequence recovery on both contact and non-contact residues on the complementary determining regions

To further understand how pretraining or fine-tuning may improve prediction for the antibody-antigen context for training strategy I, we break down the sequence recovery for the antibody and the antigen (**Figure 2B**, **SI Figure 2**). Models trained on proteins (ProtEnT, ProtPPIAbAg, ProtAbAgEnT) show higher sequence recovery on the antigen interface, whereas models fine-tuned on antibody-antigen interfaces modestly improves predictions on the antigen and significantly improves sequence recovery on the antibody interface (AbAgEnT, ProtPPIAbAg, ProtAbAgEnT).

Interestingly, while the AbAgEnT seems to perform same as or worse than the PPIEnT or PPIAbAgEnT models (**Table 1**), respectively, the breakdown in Figure 2B shows that the AbAgEnT model is perhaps specializing in highly conserved sequence patterns on the antibody interface but does not learn the diverse sequence space of epitopes that are poorly represented in the relatively small antibody-antigen datasets. Thus, pretraining on more general protein contexts (e.g., ProtAbAgEnT) is critical for prediction on the antigen interface.

To further understand the role of conserved sequence patterns in training models on antibody-antigen datasets, we calculated the sequence recovery on CDR loops. While all CDR loops undergo somatic hypermutation leading to diverse sequences that enable antibodies to recognize a wide repertoire of epitopes, five (H1, H2, L1, L2, L3) out of six CDR loops exhibit conserved sequence patterns (when structure is fixed) and antigen-induced hypervariability is limited to a few positions. Thus, breaking down sequence recovery by CDR loops and antigen-contacting residues on the antibody interface will probe the ability of different models to recover antigen-induced mutations (**Figure 2C**, **SI Figure 2**).

Pretraining on proteins (ProtAbAgEnT; dark blue bars) leads to significant improvement in sequence recovery on residues at the antibody-antigen interface across all CDR loops, including CDR H3 (**Figure 2C, SI Figure 3**). Remarkably, ProtEnT performs just as well (all residues) or better (contact-residues only) on CDR H3 as the AbAgEnT model suggesting that the sequence distribution for epitope-specific CDR H3 exhibits more general patterns and can benefit significantly from general context training. Indeed, we found that ProteinMPNN (protein-only structure conditioned sequence predictor) was remarkably good at recovering sequence of the contact residues on CDR H3 (results not shown).

#### Fine-tuning on massive antibody datasets significantly boosts overall performance on antibody-antigen sequence recovery

Following the training strategies outlined in Figure 1B, we also tested the effect of pretraining on antibody sets derived from SAbDAb and AlphaFold2 (**Figure 2D**, **SI Figure 3)**.

In **Figure 2D** and **SI Figure 3**, we show the sequence recovery for all CDR loops for training strategies II and III. First, we observed a small improvement in sequence recovery by training on larger AF2-obtained antibody training sets than the ones derived from SAbDAb alone. Remarkably, we observe that the antibody-only models (AbPlusEnT and AbEnT) show same or worse performance than ProtEnT on almost all CDR loops except CDR H1. *The inferior performance is the most pronounced for contact residues on CDR H3 where the ProtEnT is better by over 10 percentage points than AbPlusEnT and AbEnT models.* Thus, in line with our previous observation, models trained on antibody-only datasets are poor at predicting sequence of less conserved regions and contact regions.

Fine-tuning antibody-only models on the antibody-antigen datasets (AbPlusAbAgEnT or AbAbAgEnT) improves performance on all CDRs over base models (AbEnT, AbPlusEnT and AbAgEnT) on all CDRs and contact residues. The improvement is relatively small over AbAgEnT model for CDR H3 contact residues (**SI Figure 3**).

We also investigated the effect of pretraining on both proteins and antibody datasets (training strategy III; **Figure 2D**, bars in red hue). This strategy showed consistent small improvements across most CDRs (including CDR H3 and contact residues) over strategy II (pretraining on antibodies alone). Including PPI datasets during fine-tuning again slightly improved performance on contact residues on CDR H3 (**SI Figure 3**). However, no model trained with training strategies II or III matched the performance of the ProtAbAgEnT or ProtPPIAbAgEnT models that showed the highest sequence recovery across the board (except CDR H1), especially on the CDR H3 contact residues.

Here, pretraining and fine-tuning represent the physical and biological forces at play. A general model that could accurately capture how residues interact with their physical environments is not sufficient to recapitulate the antibody CDR residues due to the biological context from which these loops are derived. Thus, a model that can accurately capture both the underlying physics and the biological patterns for a given protein family is needed to design *de novo* yet native-like proteins or antigen-specific antibodies.

### Analyzing designed sequences with AlphaFold2 and language models

Next, we investigate the protein sequences sampled from ProtEnT model at 100% masking from the TS50 dataset. We are primarily interested in asking whether sequences generated from our models fold into the native structure.

**Figure 3.**
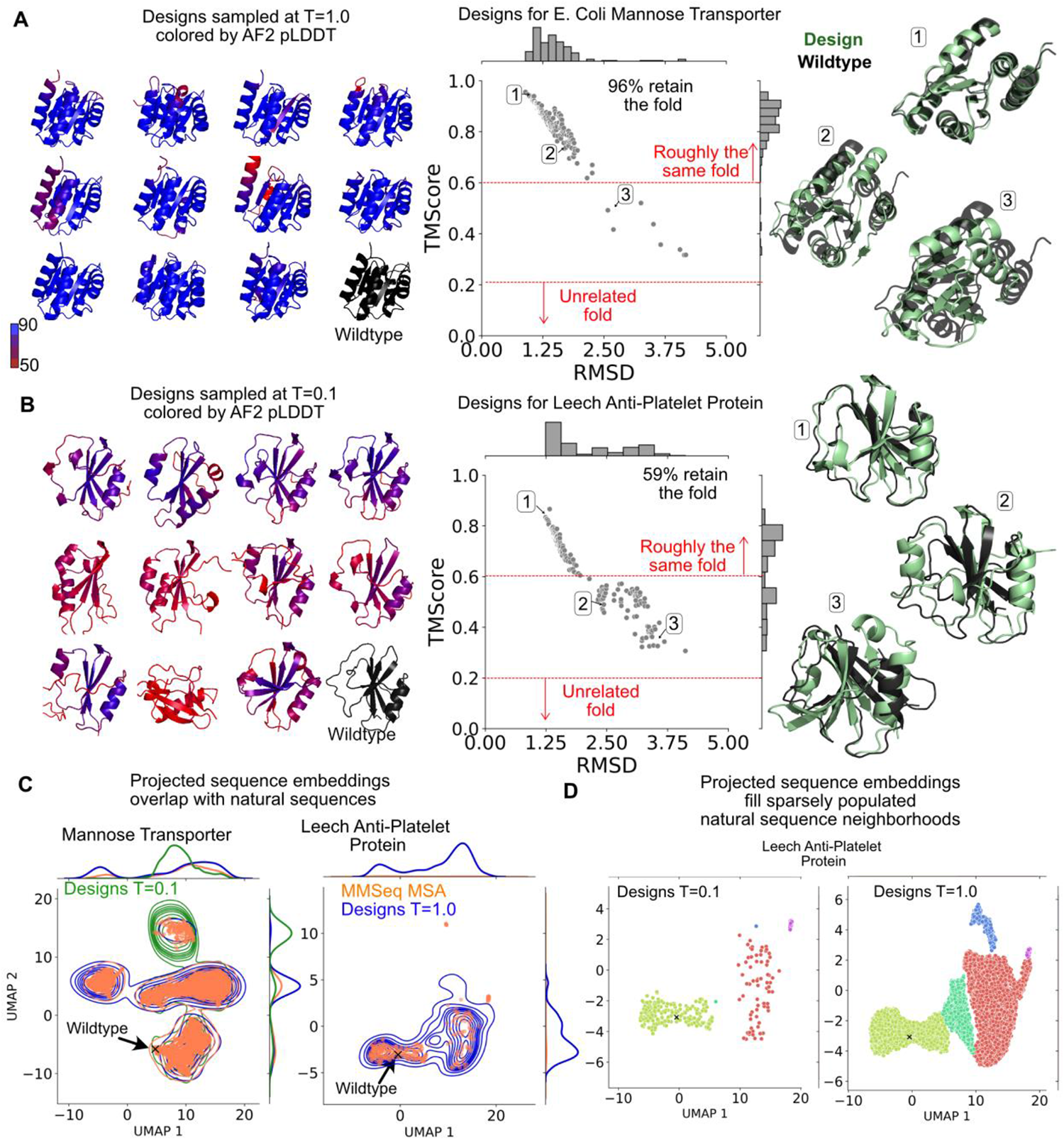
Sampled sequences generated from ProtEnT fold into target structures and access nearby sequence and structural neighborhoods. AF2 folded structures colored by pLDDT values for sequences sampled at T=1.0 for **(A)** the mannose transporter (PDB 1PDO) and **(B)** the anti-platelet leech protein (PDB 1I8N) (left), RMSD and TM score of folded structures with respect to the crystal structure (center) and three representative structures (green) selected at different TM scores aligned to the crystal structure in black (right). **(C)** Structure conditioned sequences based on the mannose transporter and anti-platelet protein sampled at temperatures 0.1, 1.0 (blue, green density plots) and 0.1 (blue density plot) respectively, in comparison with natural sequences (orange points) related to the wildtype sequence obtained with MMseq2. The sequences are embedded with ESM-1b^27^ sequence model and projected down to two-dimensions with UMAP. **(D)** Structure conditioned sequences for the anti-platelet leech protein sampled at low (0.1) and high (1.0) temperature shown as points on the projected UMAP sequence space. Sequences generated at low temperatures resemble the spread exhibited by natural sequences (C) whereas those generated at high temperatures fill regions of sequence space not populated in natural sequences. Sequences are colored by clusters assigned by DBScan^28, 29^.

#### Generated protein sequences fold into target structure and expand into nearby structural space

We obtained the maximum-likelihood designs for all 50 proteins from TS50 test set (**SI Figure 4**). Forty-three out of 50 target designs folded into a structure within an RMSD of 2 Å from the native structure (**SI Figure 4A**). Out of the seven targets for which maximum-likelihood designs failed to fold into the native structure, AF2 failed to fold the wildtype sequence of one target into its native structure (**SI Figure 4B**). For the remaining six targets, we sampled 50 designs and calculated the RMSD of the AF2 folded structure with respect to the native structure (**SI Figure 4C**). Sampled sequences resulted in significant fractions of folded structures within 2 Å RMSD of the native structure for only three out of six targets.

Among the 43 targets for which maximum-likelihood designs folded into target structures, we select two proteins for further analysis -- one with high maximum-likelihood sequence recovery ∼ 49% (Mannose transporter; PDB 1PDO) and another target with low maximum-likelihood sequence recovery ∼ 27% (Anti-platelet leech protein; PDB 1I8N).

We sampled 200 sequences conditioned on the crystal structure for each protein and folded them with AF2. For the mannose transporter, 96% of the structures retained the same fold (TM-score >= 0.5) and exhibited overall favorable pLDDT values (**Figure 3A, SI Figure 5A**). For the anti-platelet leech protein (**Figure 3B**), only 59% retain the same fold, and they exhibit lower pLDDT scores for the folded structures (**SI Figure 5B**). We also examined representative structures that fold into the target structure (higher TM scores, low RMSDs) and those that fold into structures with lower TM scores to the target structure (Selected structures 1, 2 and 3 in **Figures 3A** and **3B**). The structures with higher TM scores (“2” and “3”) exhibit many structural motifs in the starting structure with small variations such as a reorganization of α-helical arrangement in three-dimensions or change in the number of β-strands. In **SI Figure 6**, we show examples for four sampled sequences (“design”) folded with AF2 that deviate from the starting fold. To probe whether these related folds could be stabilized, we generated sequences conditioned on these sampled folds (**SI Figure 6**). Only in a few cases, we were able to find sequences that exhibited both low RMSD and AF2 structures with reasonable average pLDDT values (>=70; matching those of the wildtype sequences, **SI Figure 4B**). We also searched Foldseek^30^ to find natural proteins in the PDB and AF2^31^/ESMFold^32^ databases resembling these sampled folds and did not find any hits except for one case (**SI Figure 6**). This suggests that sequences sampled for a given structure not only fold into the target structure but also allow sampling of the structural neighborhood of the starting structure. Furthermore, low success rates of stabilizing these neighboring folds may also suggest that the fold-space of natural proteins is sparse and folds/structures that spans or connect these spaces may be unstable or only exist transiently.

#### Generated protein sequences recapitulate biologically observed sequence neighborhood of native sequence and fill out sparsely populated native sequence neighborhoods

To compare the sequence space sampled by ProtEnT and that sampled by biology, i.e., natural sequences, for our two selected targets (mannose transporter protein and anti-platelet leech protein), we sampled 10,000 sequences conditioned on their respective structures. We further searched natural sequences related to the wildtype protein with MMseq2^33^. We then embedded each sequence (sampled and MMseqs2) with ESM-1b^27^ embeddings and projected the embeddings in two dimensions as described in Hie et al^34^. The resulting projections in two dimensions reveal sequence similarity in the perspective of the protein language model in the projected space. The sampled sequences recapitulate the sequence space of the natural sequences (**Figure 3C**). Unlike the mannose-transporter protein sequence that has thousands of biologically related sequences (**Figure 3C**, left), the wildtype anti-platelet leech protein sequence has only a small number of related natural sequences and they populate a sparse region of the sequence space (**Figure 3C**, right; orange points) compared to the ProtEnT generated sequences at high temperature (blue density plot). Indeed, ProtEnT sampled sequences (**Figure 3D**) can fill out this sparsely populated sequence space (T=1.0) thereby bridging the sequence space observed in nature. *Thus, sampled sequences can expand both the local structural and sequence neighborhoods of a starting structure*.

**Figure 4.**
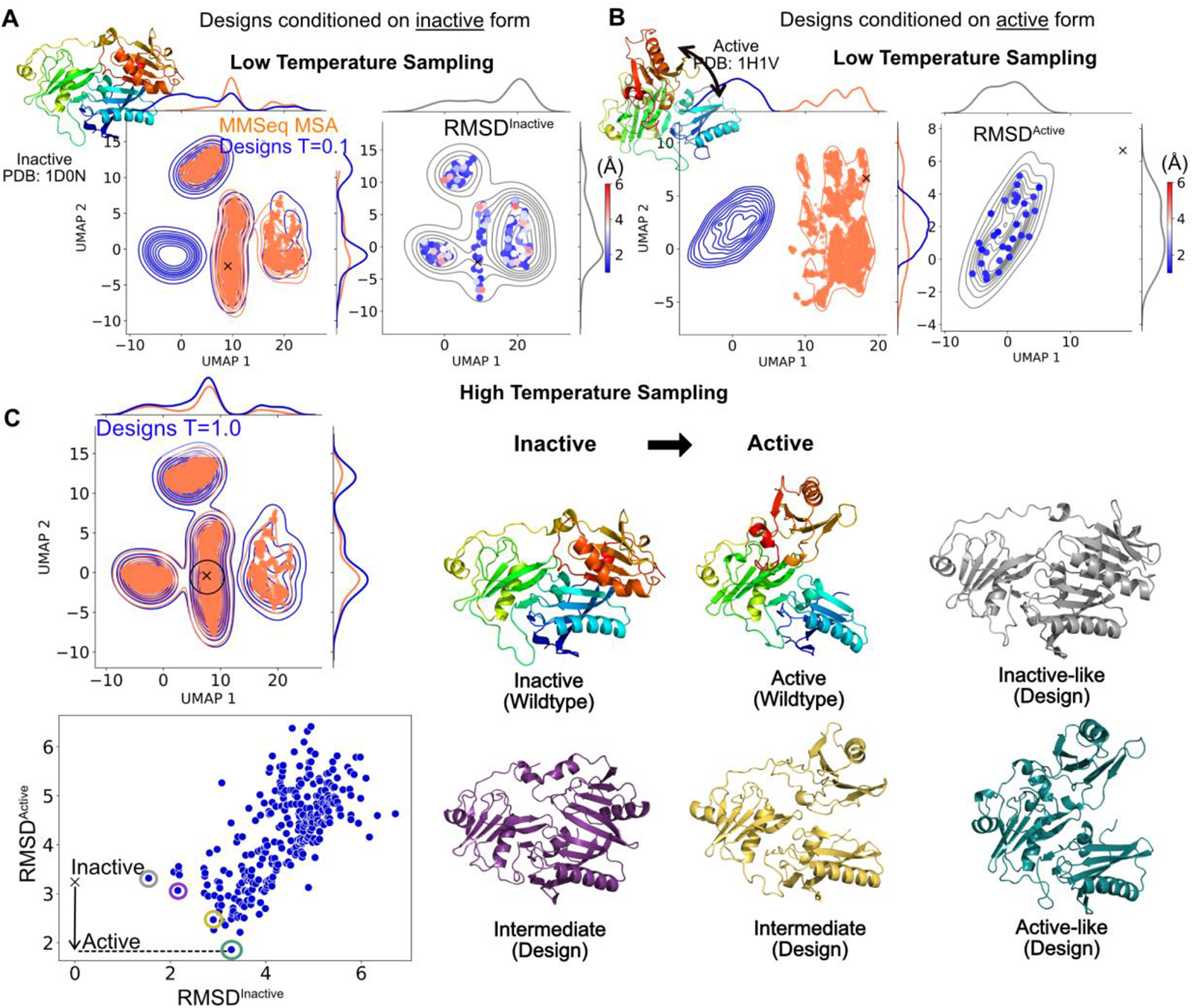
Generated protein sequences can conditionally sample structural conformers and access alternative conformations. Sequences sampled at *T*=0.1 for the *inactive* state **(A)** and *active* state **(B)** of gelsolin. (Left) Joint plot of sampled sequences (blue; shown as density contours) and natural sequences from MMseq2 (orange; shown as points) on a two-dimensional UMAP projection of ESM-1b^27^ embedded sequences. (Right) RMSD of AF2-folded structures for selected sequences generated for the inactive conformation (A) and active conformation (B), colored by the RMSD from respective conformations, on the contours of sampled sequence space (grey). **(C)** Sequences sampled at T=1.0 for the *inactive* state. (Left) See description for (A) or (B). The native or wildtype sequence is shown as a cross (x). (Bottom left) RMSD^active^ to RMSD^inactive^ for the folded AF2 structures of sampled sequences from sequences close to the wildtype sequence in the projected sequence space (black circle on the jointplot on left). (Right) Cartoon representations of the wildtype inactive (black) and active states (rainbow) and AF2 folded sequences marked with the same colors on the RMSD plot (middle). **(D)** Sequences sampled at T=0.5 for the *active* state. Descriptions for left, middle and right panels same as (C).

#### Generated protein sequences can conditionally stabilize structural conformers of the native sequence and make alternative, hard-to-sample conformations accessible

Since the models are trained to predict residue identities for a given protein structure, we asked whether the structure of conformationally plastic proteins encode flexibility and whether the sequences generated for such structures at different temperatures will reflect plasticity observed in nature. To answer this question, we chose the protein gelsolin. Gelsolin is a highly flexible protein composed of severin domains.^35^ The inactive form exhibits a closed conformation. Addition of calcium ions activates a latch-like mechanism resulting in the active open conformation (**SI Figure 7A**).

We sampled sequences conditioned on the inactive conformation (**Figure 4A**) and the active conformation (**Figure 4B**) at low temperature. At low temperatures (T=0.1), the sequences conditioned on the inactive conformation (**Figure 4A**, left) recapitulate natural sequences related to the native sequence (from MMseqs2). However, the sequences conditioned on the active conformation (**Figure 4B, left**), are distinct from natural sequences. One possible explanation for this apparent discrepancy is that the natural sequences are optimized to fold into the inactive conformation and exhibit the active conformation only when triggered by calcium ions, whereas the sampled sequences conditioned on the active conformation at low temperatures are optimal for the active conformation without the need for activation. At higher temperature (T=0.5), the sequences generated for the active conformation overlap with natural sequences (**SI Figure 7B**).

Furthermore, for each case, we folded 200 randomly sampled sequences with AF2 and calculated the RMSD’s of the structures (compare right panels for 4A and B; **SI Figure 7C**). The sequences conditioned on the inactive conformation show a wide distribution, i.e., they fold into structures that may deviate from the inactive conformation, whereas those conditioned on the active conformation exhibit very little deviation from the active conformation. This may suggest that the inactive structure is inherently more flexible than the active conformation in agreement with the biological requirement for the inactive conformation to be predisposed towards conformational change.

To investigate whether the sampled sequences could access alternative conformations at higher temperatures, we sampled sequences conditioned on the inactive conformation (**Figure 4C**). We folded 200 sequences in the neighborhood of the wildtype sequence (marked with a cross) with AF2 and calculated the RMSD of the folded structures to the active and inactive conformations (**Figure 4C**, right panel). The sampled sequences exhibit a wide range of conformations ranging from those close to the inactive conformation (grey) to the active conformation (teal) and some intermediate conformations (yellow and purple).

Thus, we can traverse the conformational space of a highly plastic protein by sampling the sequence space of the model at different temperatures and conditioning on different conformations.

**Figure 5.**
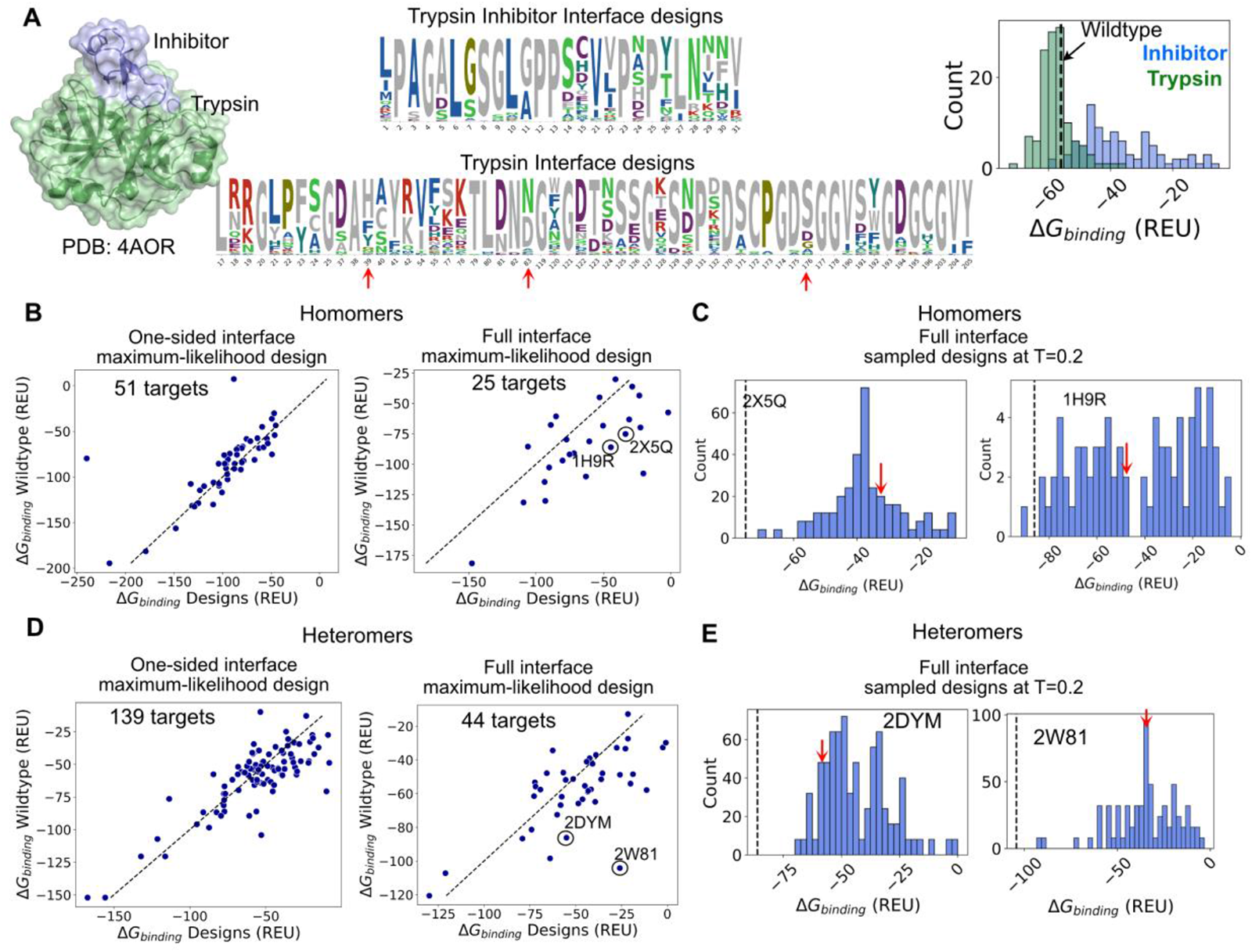
Generated interfaces recapitulate native-like binding energies (predicted by Rosetta on sequences folded with AF2-Multimer) over a wide-range of binding strengths. **(A)** (Left) Space filling model of Trypsin-Inhibitor complex. (Middle) Sequence logo for 10,000 designs sampled at T=0.5 for the interface residues on the inhibitor (top) and trypsin (bottom). (Right) *ΔG*_binding_ for designed complexes for the inhibitor (top) and trypsin (bottom). The binding energy of the wildtype complex is shown with a dashed line. (B) *ΔG*_binding_ for wildtype complexes versus *ΔG*_binding_ for sampled complexes (with ProtPPIEnT model) folded with AF2. (Left) Maximum-likelihood (T=0) sequences/designs obtained by masking interface residues for one-partner (one-sided designs) for 51 homomers from the PPI300 set. (Right) Sequences sampled by masking all interface residues for 25 homomers from PPI300 set. (C) *ΔG*_binding_ for sequences sampled at T=0.2 for two selected targets. *ΔG*_binding_ for the wildtype complex is shown with a dashed line. *ΔG*_binding_ of maximum-likelihood design is marked by a red arrow. **(D)** same as (B) for heteromeric complex. **(E)** same as (C) for targets selected from heteromeric complexes.

#### Generated interfaces retain binding with partner and expand the range of binding affinities for a given interface

We next asked whether sequences sampled in the context of their binding partner retained binding and whether the binding strengths recapitulated those of the native binders. To this end, we generated sequences by masking interface residues in the context of the partner by folding with AF2-multimer followed by calculating the free-energy of binding (*ΔG*_binding_) with Rosetta’s interface analyzer^36^.

First, we considered the trypsin-trypsin inhibitor complex, which was the target with the highest sequence recovery ∼ 60%, from the PPI300 test set (**Figure 5A**). We sampled 150 sequences at *T* = 0.5 for the trypsin enzyme with a fully masked interface in the context of the unmasked inhibitor and vice-versa; the resulting sequence logos are shown in Figure 5A (native residues are shown in grey). We folded the resulting sequences with AF2 multimer and characterized the *ΔG*_binding_ with Rosetta’s interface analyzer also shown in **Figure 5A**. For the trypsin designs, the majority of sampled sequences exhibit binding energies the same as or better than the wildtype complex. For the inhibitor designs, most of the sampled sequences exhibit worse binding energies than the wildtype complex, with a few sampled sequences exhibiting a slightly better binding energy than the wildtype complex. This is not surprising as the catalytic pocket in trypsin evolved for catalysis whereas the inhibitor evolved for binding the catalytic pocket. It is, therefore, possible to optimize trypsin’s catalytic pocket for binding at the cost of catalytic activity but challenging to further improve the inhibitor for binding. As expected, a significant percentage of designed sequences remove catalytic residues (**Figure 5A**; red arrows) though many designs also retain the wildtype residues. Surprisingly, the design with the highest binding energy (as estimated by Rosetta) retains the catalytic triad but loses a cysteine (at position 40 in **Figure 5A**; position 64 in PDB 4AOR) that forms a disulfide bond with another cysteine that probably provides additional rigidity to the catalytic triad not essential for binding.

#### Maximum likelihood interface designs match native free-energies of binding over a wide-range of interfaces and binding strengths

We further investigated whether designed interfaces recapitulate binding strengths of native binders (as we did for the trypsin-inhibitor complex) over a large number of targets. We asked whether the coarse-grained structural-context of the binding partner is sufficient (coarse-grained structure alone) to obtain native-like binding strengths or whether both the shape and chemical composition of the binding surface (sequence with structure) are essential for generating binders with native-like binding strengths. To answer these questions, we carried out two sets of interface designs -- one-sided and full interface designs. We calculated the binding energies of designed interfaces by redesigning a subset of the PPI300 test set and selecting the maximum-likelihood designs (argmax over predicted amino-acid likelihoods) for each target (**Figure 5B, C, D, E**).

For one-sided designs, we fixed one partner and obtained the maximum-likelihood interface design for the other partner for 102 homomeric targets (102 complex designs), folded the resulting complexes with AF2-multimer, identified targets for which the folded structures were within 2.0 Å of the target structure and calculated their binding free energies with Rosetta and compared them to the binding free-energy of the wildtype complex. Only 51 of 102 designs folded into structures within an RMSD of 2.0 Å from the target complex structure (**SI Figure 8A**). Remarkably, even though the designs exhibit about 40% median sequence identity with the wildtype sequence, the binding strengths of the designed complexes (as estimated by Rosetta) recapitulate those of the wildtype complexes for a wide range of binding free energies (**Figure 5B** left panel).

For the full-interface designs, we masked the interface residues for both partners at the same time for 25 homomeric targets (25 complex designs). For the 21 designs that folded into structures with RMSD under 2.0 Å (**SI Figure 8B**) from the target complex structures, we obtained the binding free energies with Rosetta for the maximum-likelihood designs as described above (**Figure 5B**, right panel). In this case, while there is a strong correlation between the binding energies of the designs and the wildtype complexes, designs exhibit worse binding energies than the wildtype. This is not surprising as the designed interfaces only see the structural context of the partner but lack the sequence context (as it is masked, and all residue identities are chosen simultaneously from the maximum likelihoods).

We reasoned that better interface designs could be found by sampling many sequences from the model for a particular target. For two targets (2X5Q, 1H9R) whose designs had weaker binding energies than the wildtype (Figure 5B right panel), we sampled 200 sequences at T=0.2 and repeated the analysis (**Figure 5C**). As we suspected, the binding energies of the sampled sequences (bars) result in designs close to the wildtype binding energy (dashed line) and the maximum likelihood design (red arrow).

We repeated the above analysis for 100 targets (or 200 one-sided designs) and 100 (full-interface design) heteromeric targets from the PPI300 test set (**Figures 5D, E**). For two out of 100 targets, AF2 failed to fold the wildtype sequences within 2 Å RMSD of the native structure. These were omitted from further analysis. For one-sided designs, 139 of 196 folded into complex structures with RMSD < 2.0 Å with respect to the wildtype (**SI Figure 8C**). For these 139 designs, we compare the binding energies to their respective native sequences. For the full interface design for the heteromeric targets, only 44 out of 96 maximum-likelihood designs folded into structures with RMSD under 2.0 Å from the wildtype complex (**SI Figure 8D**). For both one-sided and full interface design, we observe larger deviations from the wildtype binding energies (**Figure 5C, D**). To further investigate designs with significantly weaker binding energies than the wildtype, we sampled 200 sequences for two targets (2DYM, 2W81; **Figure 5E**) using a finite temperature. Sampling more sequences improves binding free energies for full interface designs over maximum likelihood designs (red arrows).

From the above analysis we conclude that one-sided interface designs that folded into the target structure recapitulate wildtype-like binding energies for a wide range of binding strengths. Furthermore, we also find that designing optimal interfaces requires both structure (shape) and sequence (chemical composition) context of the partner. For the full-interface design, with our one-shot decoding strategy, the sequence context of the partner is absent leading to suboptimal designs that are inferior in binding strengths to the wildtype complex. A sequence decoder (like ProteinMPNN or IF1) may be able to overcome this weakness.

**Figure 6.**
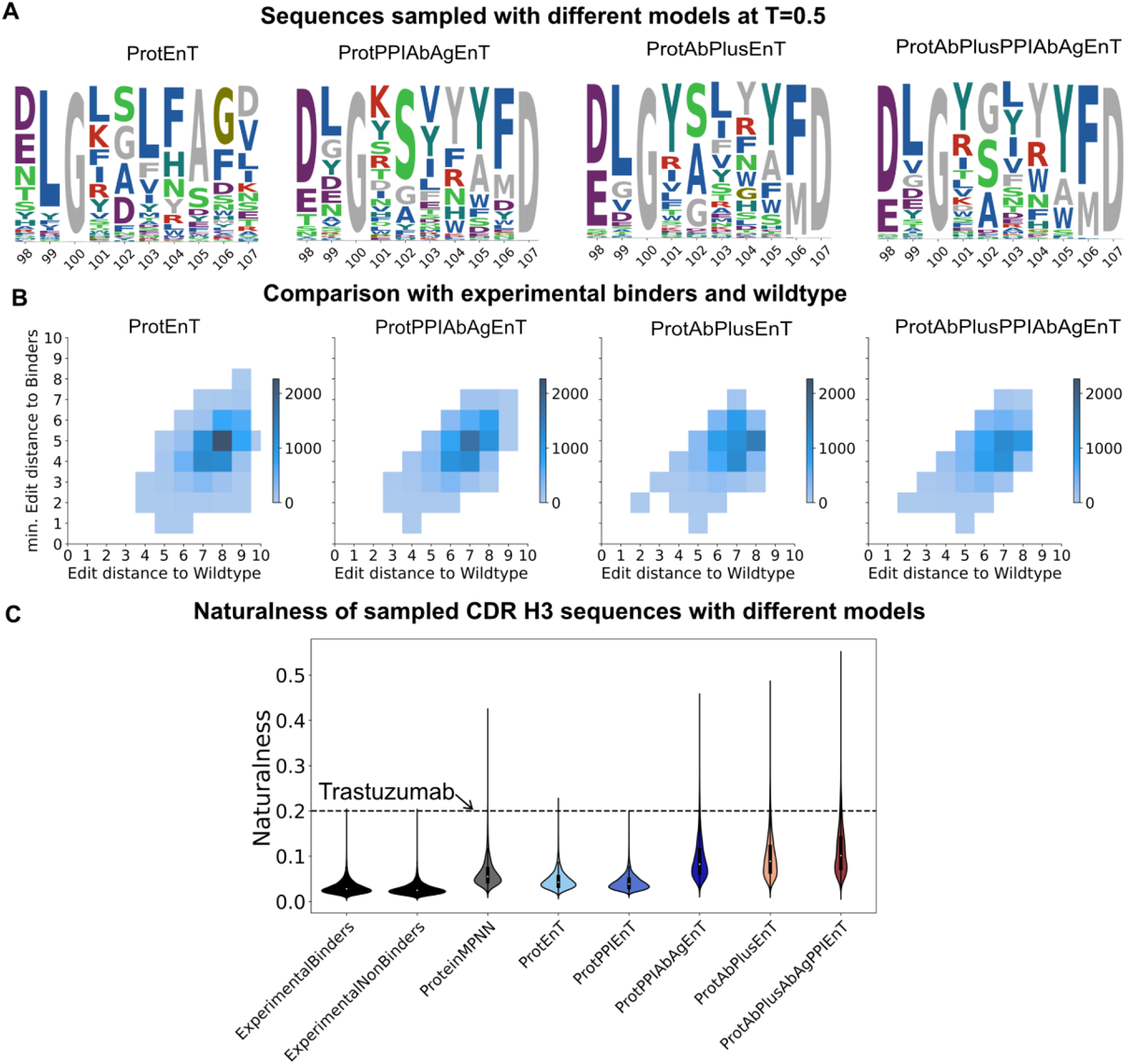
Evaluation of sampled sequences for CDR H3 for the Trastuzumab-HER2 antibody antigen complex. **(A)** Sequence logos for 10,000 sequences sampled at *T* = 0.5 for each model for 10 positions on CDR H3 of the trastuzumab antibody conditioned on the trastuzumab-HER2 complex structure (PDB 1N8Z^37^). **(B)** Edit distance of sampled sequences from the wildtype trastuzumab sequence (x-axis) and minimum edit distance of each sequence from the set of sequences of experimental binders (y-axis) for four models. **(C)** Naturalness of 10,000 CDR H3 sequences conditioned on trastuzumab-HER2 complex sampled from models presented in this work, with ProteinMPNN^14^, experimental binders and non-binders from Mason et al.^38^, and the wildtype Trastuzumab antibody.

#### Generated sequences from antibody-specific models resemble natural antibody sequences and recapitulate sequence preferences observed in experimentally screened CDR libraries

We could not analyze antibody designs in the way we analyzed designs for protein complexes in the previous section because there is still no fast and robust machine learning model or *in silico* method to obtain accurate structures of antibody-antigen complexes from sequences. Thus, to compare sequences sampled for antibody CDR loops from our models, we considered the Trastuzumab-HER2 antibody-antigen complex (PDB 1N8Z^37^). For this antibody-antigen complex, Mason et al.^38^ have experimentally obtained 11,000 unique CDR H3 sequences for the Trastuzumab antibody that retain binding (after multiple rounds of screening for expression and binding) to HER2.

We sampled 10,000 antibody sequences with different models conditioned on the crystal structure at *T* = 0.5 (**Figure 6A**). Comparing sequence distributions from models that were trained with (ProtAbPlusEnT, ProtAbPlusAbAgEnT) and without (ProtEnT, ProtPPIAbAgEnT) large antibody datasets, we observe a few key differences. Out of the four models shown, only ProtEnT, which was trained without any antibody information, fails to recover Y as the most preferred amino acid at position 104 – a residue found to be critical for retaining binding to the antigen in single-site mutagenesis experiments. All models show strong preference for amino acid residues G, S and A at position 102, another position found to be important for binding in experiments and structure-conditioned design studies^39^. In line with our results on sequence recovery (**Figure 2**), the model fine-tuned with training strategy I (ProtPPIAbAgEnT model) best recapitulates both conservation patterns in antibodies and non-conserved antigen-driven mutations.

We also calculated the edit distance (Levenshtein distance) between each sampled sequence and minimum distance to a binder from the experimental set (**Figure 6B**). Remarkably, all four models sample sequences with up to 90% sequence identity with an experimental binder (min. edit distance of one) and up to 60-80% sequence identity with the wildtype sequence (edit distance of four to two).

Finally, we calculate the “naturalness” with AntiBERTy^40^ (a masked language model) of the CDR H3 sequences sampled from different models. We also calculated the naturalness of the wildtype Trastuzumab CDR H3, sequences sampled with ProteinMPNN (*T* = 0.5), and experimental binders and non-binders from the Mason et al.^38^ set (**Figure 6C**). Experimental binders’ sequences, which were obtained by exhaustively scanning the single-site sequence space of CDR H3 followed by combinatorial library design, exhibit much lower naturalness than Trastuzumab sequence. This suggests that naturalness itself is not important for retaining binding to the target. When comparing different models, it is evident that protein-only models generate sequences with less naturalness than models trained on antibody datasets. Thus, fine-tuning on antibody datasets leads to more “natural” sequences.

## Conclusions

In this work, we present a masked label prediction model for proteins that learns in the context of the sequence and structure of the unmasked residues. We further investigated whether such contextual encodings can be fine-tuned for specialized contexts such as the antibody-antigen interface.

Even though the models were trained at a low mask rate (15 – 25 %), at 100% masking during inference, the model trained on protein datasets (ProtEnT) recovers 37% of the native sequence suggesting that over one-third of the amino acid label may be determined from its structural context alone without any autoregressive decoding of the sequence. Other recent works such as PiFold have demonstrated that deep sequence and structural encoders can improve performance on inverse folding tasks with one-shot decoding.^15^ Moreover, while representation learning on sequences has been widely adapted to different tasks and protein domains, representation learning on structures and structure-sequence contexts is relatively nascent.^7–9^ Our work demonstrates that joint structure and sequence representation learning can be applied to boost performance on data-sparse regimes such as the antibody-antigen complexes. In line with a recent study on evaluating the available data on antibody-antigen complexes and affinity,^41^ our work demonstrates that the diversity and sequence-structure space coverage in experimental antibody-antigen datasets is insufficient to train robust models for binding or affinity prediction. Transfer learning from data-rich regimes offers one way to alleviate this problem.

We also show that sequences sampled from the masked model fold into target structures, although the AF2 model confidence (as measured by pLDDT of the AF2 folded structures) deteriorates for targets with lower native sequence recovery. The generated sequences recapitulate the sequence space of the sequence neighbors of the wildtype sequence, explore local structural neighborhoods and, for structures known to exhibit high plasticity, make hard-to-access, alternative conformations accessible. Recent works such as ESM2^32^ and ESM-IF1^26^ have also demonstrated the ability of deep learning models to extrapolate sequence and structural space beyond natural proteins and recapitulate sequence likelihoods of plastic proteins.

We also explore the properties of residue designs at protein-protein interfaces and find that maximum-likelihood designs (one-sided) from masked models recapitulate native-like binding energies for a wide-range of binding strengths. However, the binding energies of the maximum-likelihood interfaces sampled without both sequence and structural context of the partner (i.e. when interface residues on both partners are masked) are significantly lower than those of the wildtype complex.

We find that pretraining improves native sequence recovery in specialized contexts, especially for hypervariable contact residues on antibody CDR loops, and that fine-tuning on antibodies improves the “naturalness” of generated antibody sequences. Similar results have been reported for antibody-specific infilling language models such as IgLM.^42^ By developing models for specialized contexts such as soluble proteins (SolubleMPNN), an extension of the inverse folding model ProteinMPNN trained on soluble proteins only, Goverde et al. successfully designed soluble versions of membrane proteins.^43^

Our approach sheds light on transferable patterns within proteins, between proteins and protein-protein interfaces and from proteins or protein interfaces or antibodies to antibody-antigen interfaces. Our work also highlights the trade-off between learning sequence/structural patterns specific to a protein family such as antibodies and learning distribution of the three-dimensional patterns of protein residues in physical environments. Further work on understanding specific structural and sequence patterns in binding surfaces, especially those in specialized contexts such as the paratope of antibodies could pave the way for *de novo* design of antibodies.^39, 44–47^

We hope that future work can shed light on the different choices of model, representation and architecture -- deeper sequence encoders, multiscale representations (coarse-grained to fine-grained), masking schemes, different pretraining and fine-tuning schemes and the use of a pretrained and/or context-specific sequence decoder for sequence generation.

## Methods

### EGNN

Satorras *et. al.*^17^ proposed an equivariant graph neural network (EGNN) for graph structured data. For a graph 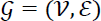 with nodes 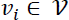 and edges 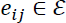. Each node has a coordinate *x_i_* ∈ ℝ^3^ associated with it. The operations in a single EGNN layer are defined as follows:

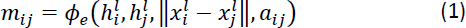

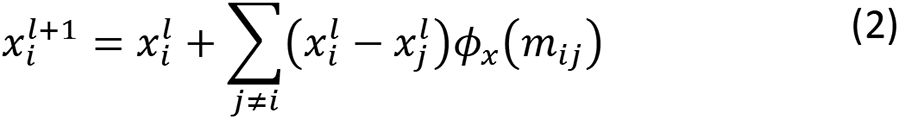

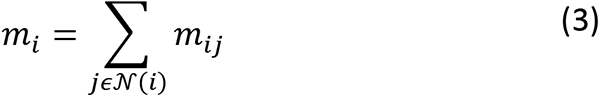

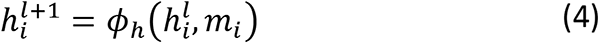

where 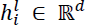 is the *d*-dimensional embedding of the node v_*i*_ at layer *l*. a_*i,j*_ are the edges *i.e.,* 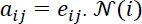 represents the set of neighbors of node *v_i_*. *ϕ*_*e*_, *ϕ*_*h*_ and *ϕ*_*x*_ are the edge and node operations, which are approximated as multilayer perceptrons in the EGNN^17^.

### E(n)Transformer

The E(n)Transformer, to the best of our knowledge, is a variation on the EGNN, proposed by Phil Wang (https://github.com/lucidrains/En-transformer). In the E(n)Transformer, attention is introduced to update equations (1), (3) and (4) as follows:

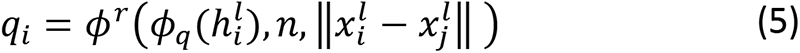

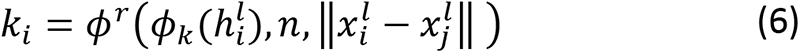

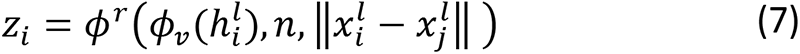

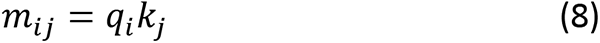

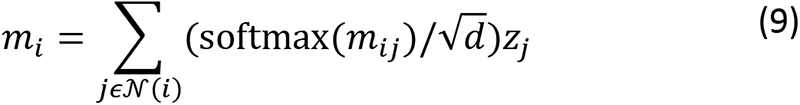

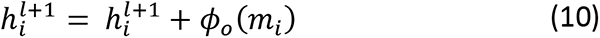

where 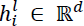 is the *d*-dimensional embedding of the node ν_*i*_ for *one attention head* at layer *l*. Ν(*i*) represents the set of neighbors of node 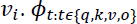 are learned linear projections to produce queries, keys, values, and outputs^25^. Equations 6-8 replace the edge embeddings *m_ij_* with attention-based query-key-value relationships. Since the formulation only depends on the relative distance, equations 6-8 are invariant to rotations and translations. *ϕ*^*r*^ are rotary positional embeddings to combine/inject amino acid residue position “n” and relative distances with query, key and value embeddings.^48^

And lastly, node embedding updates in original formulation (equation 4) now have a residual component (equation 10) same as the transformer^25^. Additionally multiple heads are employed and equations 5-10 are applied for each head. The “talking-heads attention” formulation is used to aggregate information from all the heads.^49^

### Model Architecture

The model consists of two linear layers that project node and edge features respectively from input dimensions to the dimension of the hidden layers. The projection layers are followed by 4 E(n)Transformer layers, with a hidden dimension of 256 and 8 attention heads. The output from the last E(n)Transformer layer is reshaped from a per-atom representation to a per-residue representation followed by a linear layer that projects down from the hidden dimension to 20, the number of amino acid labels.

### Datasets

#### Protein Test Sets

| Test Set Name | Description | Task | Size |
| --- | --- | --- | --- |
| **ProteinTS50** | Set of 50 proteins<sup>50</sup> | Sequence Recovery | 50 |

#### PPI Test Sets

| Name | Description | Task | Size |
| --- | --- | --- | --- |
| **PPI300** | Subset of MaSIF-search<sup>20</sup> test set without antibody-antigen pairs (Full list provided as a csv file in Supplementary material.) | Sequence Recovery | 302 |

#### Antibody-Antigen Test Sets

| Name | Source | Description | Task | Size |
| --- | --- | --- | --- | --- |
| **AbAg70** | RAbD Dataset<sup>51</sup> | Curated dataset of solved structures of variable domain of conventional antibodies in complex with their antigens | Sequence Recovery | 60 complexes |
|  | vHH Dataset (introduced in this work; SI Table 1.) | Solved structures of variable domain of single domain antibodies in complex with their antigens | Sequence Recovery | 10 complexes |
|  | Two additional antibodies (1MLC<sup>52</sup> - D44 antibody, 1N8Z<sup>37</sup> - Trastuzumab) | Solved structures of variable domain of two commonly benchmarked antibodies in complex with their antigens | Sequence Recovery | 2 complexes |
| Trastuzumab-HER2 CDR H3 set | Mason et al.<sup>38</sup> | Combinatorial CDR H3 library design followed by multiple rounds of screening for expression and affinity | Overlap with H3 variants, naturalness | 11,300 (binders)<br>27,000 (non-binders) |

#### Protein Train Set

For training on proteins, we use protein chains curated in the “Sidechainnet” datasets for CASP12 at 50% thinning^18, 19^. To remove chains with high sequence identity with proteins, complexes and antibodies or antibody-antigen complexes in the test sets, we remove all proteins in the training and validation sets with more than 40% sequence identity with any non-antibody protein chain in the test sets. For antibodies, we remove all chains in the protein train and test sets with more than 70% sequence identity with the H or L chains in the antibody test sets using the CDHIT server.^53–55^

#### PPI Train Set

For training on protein-protein interfaces, we used the non-redundant heterodimer protein-protein interface dataset of about 5,500 complexes from MaSIF.^20^ The training and test sets were generated by a hierarchical clustering of the structures according to the TM-align score at the interface using scikit-learn’s hierarchical clustering.^20^ We additionally removed all antibody-antigen and nanobody-antigen complexes from this dataset (by cross-checking PDB ids with the SAbDAb^21^ database) resulting in about 4100 complexes. About 300 of 4100 complexes were reserved as the test set while the rest were used for training. We further removed all chains with more than 40% sequence identity to the antigen chains in the antibody-antigen datasets with the CDHIT server.

#### Antibody-Antigen Train Set

For training on antibody-antigen interfaces, we obtained all antibody-antigen interfaces from the SAbDAb database^21^ (February, 2022). We removed all interfaces with more than 40% sequence identity with the non-antibody chains in the antibody-antigen and protein-protein test sets and all antibody (H or L) chains with more than 90% sequence identity with the antibody chains in the antibody-antigen test set.

#### Antibody Train Set

For training on the variable region of antibodies, we obtained all high-resolution antibodies and nanobodies (≤ 3.5 Å) from the SAbDAb database^21^ (February, 2022). We additionally used a set of 40,000 unpaired and unpaired antibody structures generated with AlphaFold2 from a previous study.^56^ Antibodies with more than 90% sequence identity to any chain in the antibody-antigen test set were removed.

### Dataset Preparation

For the protein dataset, to fit a full protein on a GPU, we crop proteins with lengths greater than 350 residues. The crops are chosen arbitrarily at runtime. For protein-protein datasets, we use cropped structures by including all interface residues (C_β_ atoms within 10 Å of any residue on the binding partner) and residues neighboring interface residues (C_β_ atoms within 10 Å of any residue on the binding partner) to provide additional context for the contact residues. For antibody-antigen interfaces, we retained all residues on the variable domain and cropped only the antigen.

### Training and Fine-tuning

During training, for a small percentage (15% – 25%) of the residues per training sample (protein), we corrupt/mask the amino acid identity and the C_β_ coordinates. The model is trained to predict the correct amino acid identity of these masked residues with a categorical cross entropy loss for the amino acid identity prediction.

For fine-tuning, we use the weights from the parent/baseline model (warm start) as the starting point for training the fine-tuned model. Besides training on the fine-tuning dataset, we also use the original dataset that was used to train the parent/baseline model. Since the parent model datasets are usually larger than the fine-tuning datasets, at each epoch, we sub-sample from the parent model datasets (example protein dataset) the same number of samples available in the fine-tuning dataset (example PPI dataset).

For training on proteins or antibodies, residues for masking are chosen randomly at runtime. For protein-protein and antibody-antigen training, only contact residues are masked (15%) since these residues include the full context of the neighboring residues. We mask non-contact residues at a much lower rate (5%) to introduce some noise in prediction of the contact residues.

### One-shot sequence decoding and sampling sequences from models

For each masked or design position *i* on the protein, the last layer of the model predicts the likelihood of each of the 20 amino acid residues *j* as *P_i,j_*. To sample sequences at a given temperature (*T*), we normalize the likelihoods from the model at the given temperature with a softmax operation as follows:

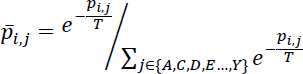

To obtain the designs, we sample from the resulting distribution independently at each design/masked position. For maximum-likelihood designs, we simply take the argmax (amino acid residue with highest likelihood) at each design/masked position over the amino-acid predictions *p_i,j_*.

### Sequence Projection with ESM

We applied the evovelocity framework developed by Hie et al.^34^. We model sequence similarity between pairs as the Euclidean distance between their ESM-1b^27^ (1 x 1280 dimensional) embeddings. The sequences are represented by their ESM embeddings and the embeddings are projected down to a 2-dimensional space with UMAP.

### Folding sequences into structure

We folded sequences for single proteins and complexes with the AlphaFold2 implementation on Colabfold^57^ (December 2022) following instructions on https://github.com/sokrypton/ColabFold and with IgFold^22^ following instructions on https://github.com/Graylab/IgFold.

### Calculation of binding free energy with Rosetta

The free energy of binding was calculated using the InterfaceAnalyzer application in PyRosetta^36^. For each sampled sequence, we obtained the structure with AF2-multimer with Colabfold, and then we relaxed the interface with Rosetta with five decoys and selected the decoy with the lowest free energy of binding (reported by InterfaceAnalyzer as dG_separated).

### Naturalness calculation with AntiBERTY

Following the definition by Salazar et al.^58^ for masked language models, Bachas et al.^59^ defined the naturalness (n_s_) of an antibody sequence as the inverse of its pseudo-perplexity . Salazar et al.^58^ define the pseudo perplexity (PPPL) as a function of the pseudo log-likelihood (PLL) of a masked language model. To estimate the pseudo-perplexity, we use the masked language model AntiBERTy^40^ that is trained on antibody sequences from the Observed Antibody Space^60, 61^.

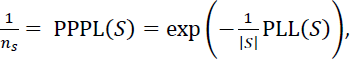

where S is the protein sequence and |S| is its length.

For naturalness reported in Figure 6, we aggregated naturalness over the 10 designed positions on CDR H3 (95 – 101 in Chothia numbering^62^).

## Code and Data Availability

The code will be available on Github at https://github.com/GrayLab/MaskedProteinEnT. Trained models, sampled sequences and results for all test sets reported in the work will be deposited on Zenodo.

## Funding

This work was supported by NIH grants R01-GM078221 and R35-GM141881. J.A.R was also supported by the Johns Hopkins-AstraZeneca Scholars Program.

## Acknowledgements

We thank Dr. Jeremias Sulam (JHU) for helpful suggestions at the early stages of the project. We especially thank former and current Graylab members, namely, Dr. Rahel Frick, Lee Shin Chu, and Dr. Ameya Harmalkar for helpful discussions. We acknowledge high performance computing resources from MARCC, ARCH and XSEDE. A preprint of this work is available at BIORXIV.

## Author contributions

S.P.M, J.A.R, J.J.G conceptualized the work

S.P.M, wrote the code

S.P.M wrote the original manuscript

S.P.M wrote analysis scripts, tested the code, and analyzed data

S.P.M, J.A.R, J.J.G interpreted data and edited the manuscript

J.J.G acquired the funding and supervised the project

## Competing interests

The authors declare no competing interests.

## Notes

### Competing Interest Statement

The authors have declared no competing interest.

### Summary of Updates

Missing reference to Phil Wang's EnTransformer added. Added "antibody" to the title.

