## Supplementary Figures and Tables for "Contextual protein and antibody encodings from equivariant graph transformers"

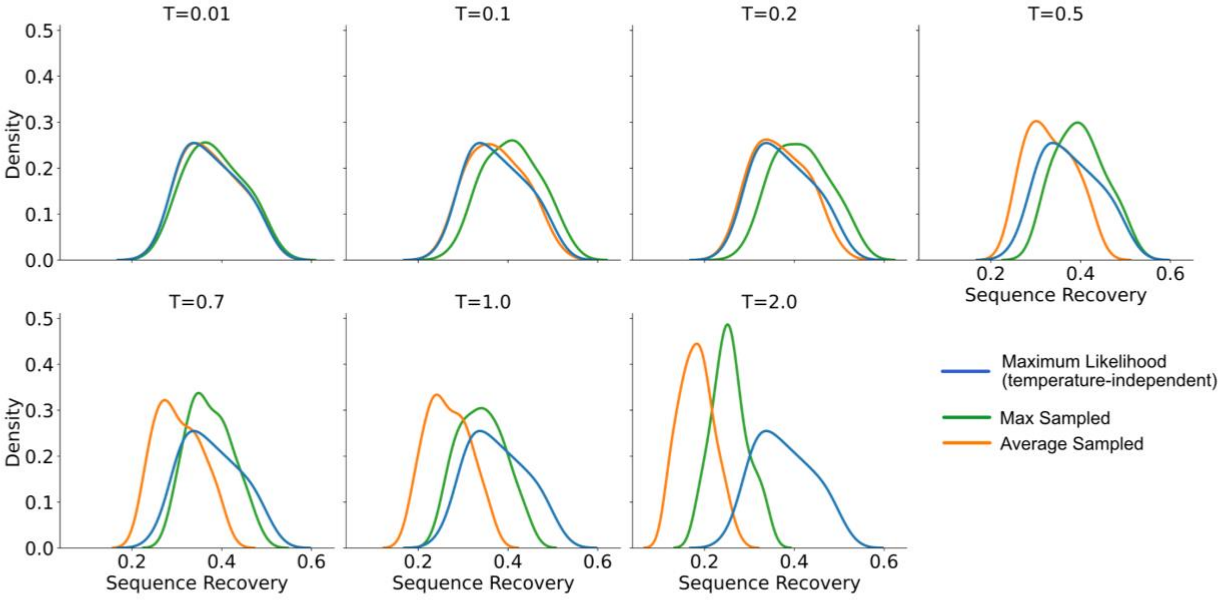


**SI Figure 1. Sequence recovery on the TS50 test set as a function of different masking rates and temperatures at 100% masking. At low temperatures, the average (average over 100 samples), maximum-sampled (maximum sequence recovery among 100 samples), and maximum-likelihood (T=0) or argmax sequence recovery overlaps. At intermediate temperatures (0.1 – 0.7), the maximum-sampled recovery is higher than maximum-likelihood and average recovery and at higher temperatures, both average recovery and maximum-sampled recovery are significantly lower than maximum-likelihood recoveries.**

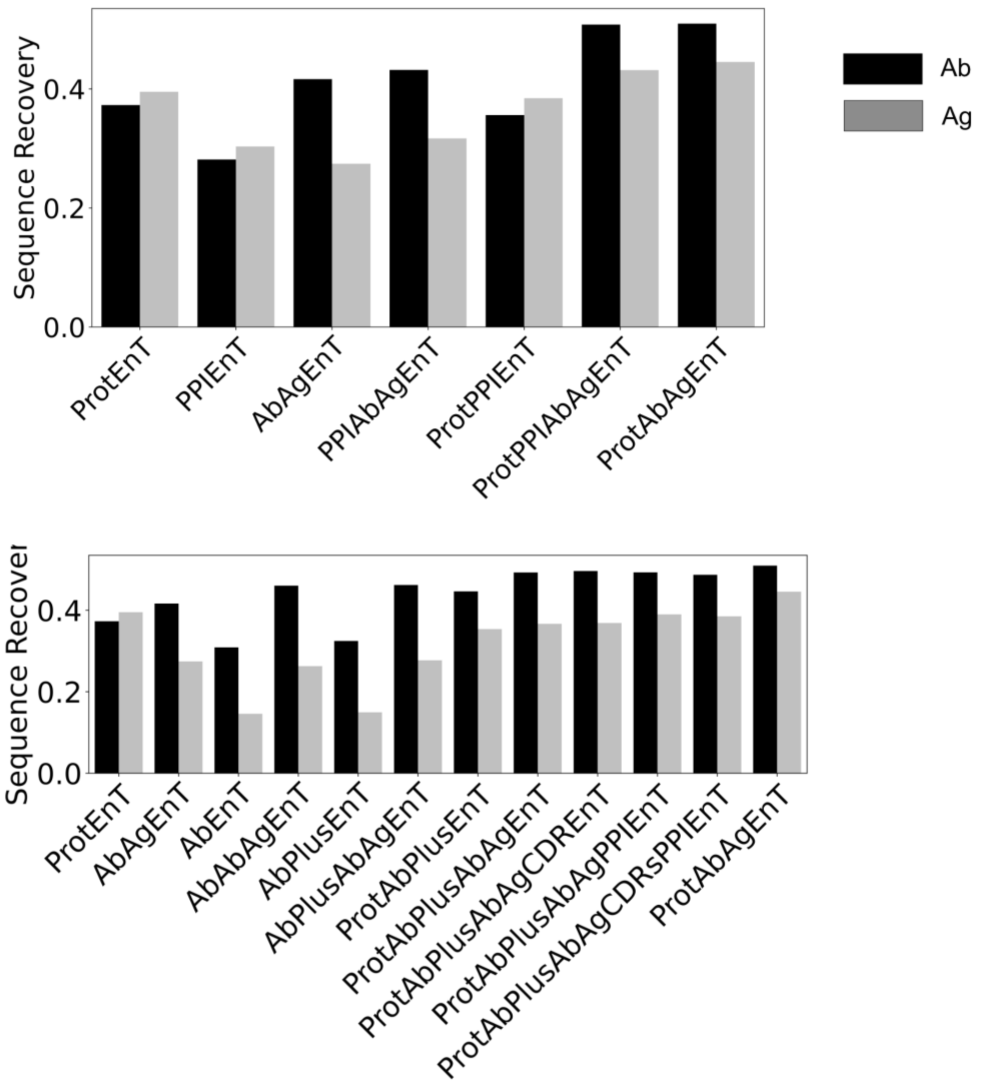


**SI Figure 2. Sequence recovery for the AbAg70 test set for all models from training strategy I (Top) and training strategies II and III (Bottom). We include ProtAbAgEnT model from training strategy I in the bottom panel for side-by-side comparison. The reported recovery is the average recovery on the test set over 100 sequences sampled per each target.**

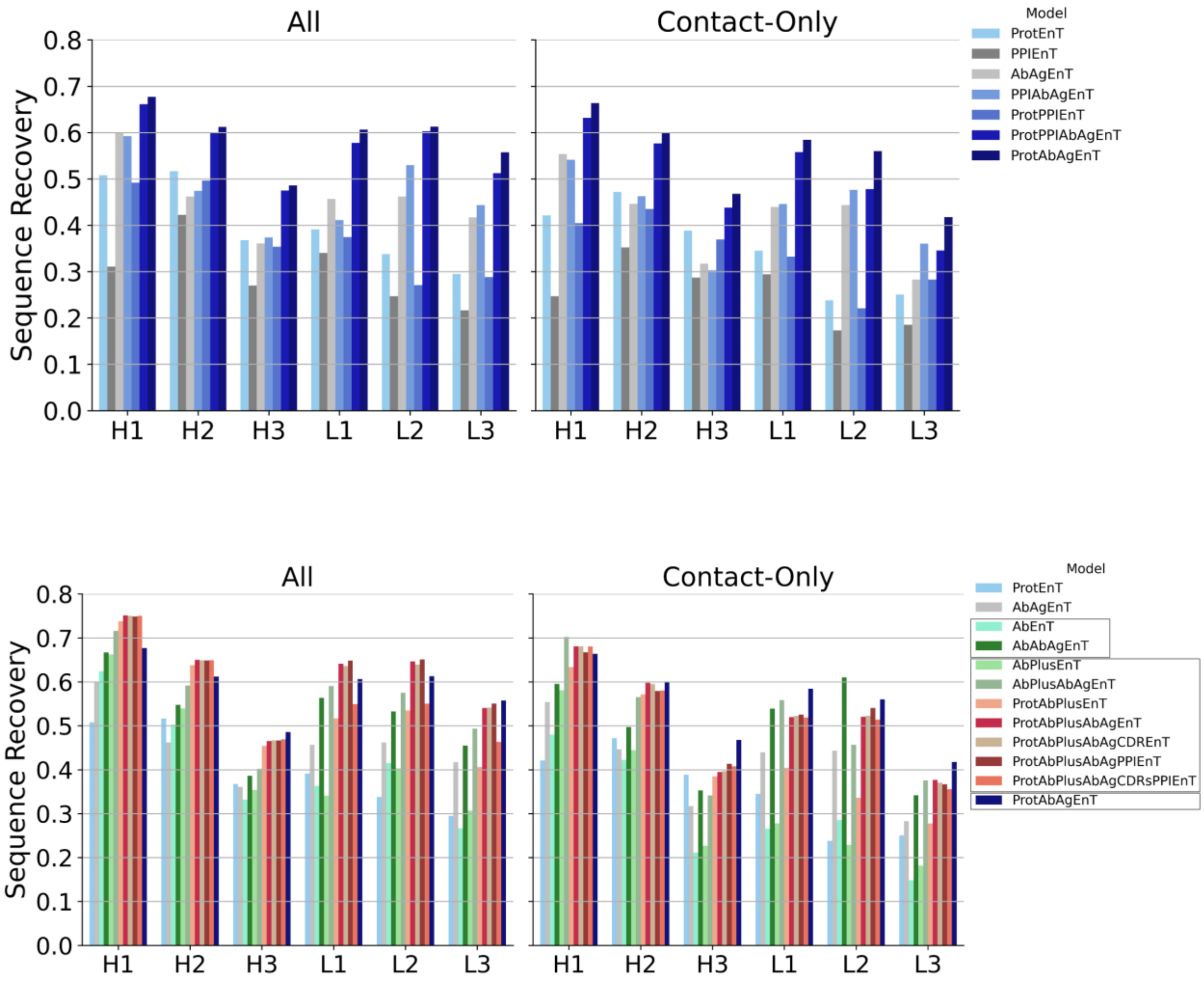


**SI Figure 3. Per-region sequence recovery for training strategies I (Top), II and III (Bottom).** Sequence recovery on the AbAg70 test set for either a masked CDR loop (left subplot) or only masked contact residues on the CDR loop (right subplot). For left subplots, we report median sampled recovery over 100 sampled sequences for each target. For the right subplot, since the number of contact residues can vary significantly per loop per target, we report an average recovery over the full dataset (i.e. (total matches with native sequence) / (total contact residues sampled per loop)).

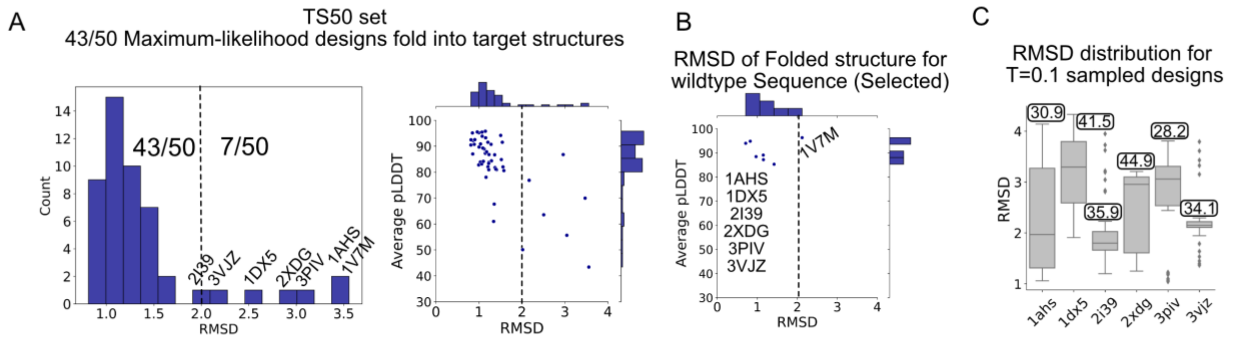


**SI Figure 4. AF2 folding metrics for proteins from TS50 test set. (A**) (Left) Distribution of RMSD (Å) and (Right) average pLDDT from AF2 versus RMSD (Å) for the maximum likelihood designs. Seven designs with RMSDs >= 2 Å are labeled. **(B)** RMSD (Å) of AF2 folded wildtype sequences for comparison. Only one target (1V7M) AF2 structure has an RMSD >= 2.0 Å. **(C)** RMSD distribution of 50 sampled designs for six targets.

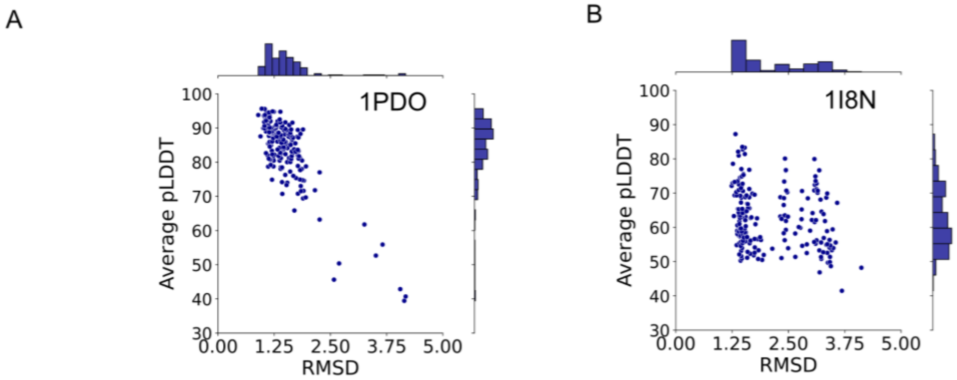


**SI Figure 5. pLDDT of 200 sampled and AF2 folded sequences for (A)** 1PDO and **(B)** 1I8N.

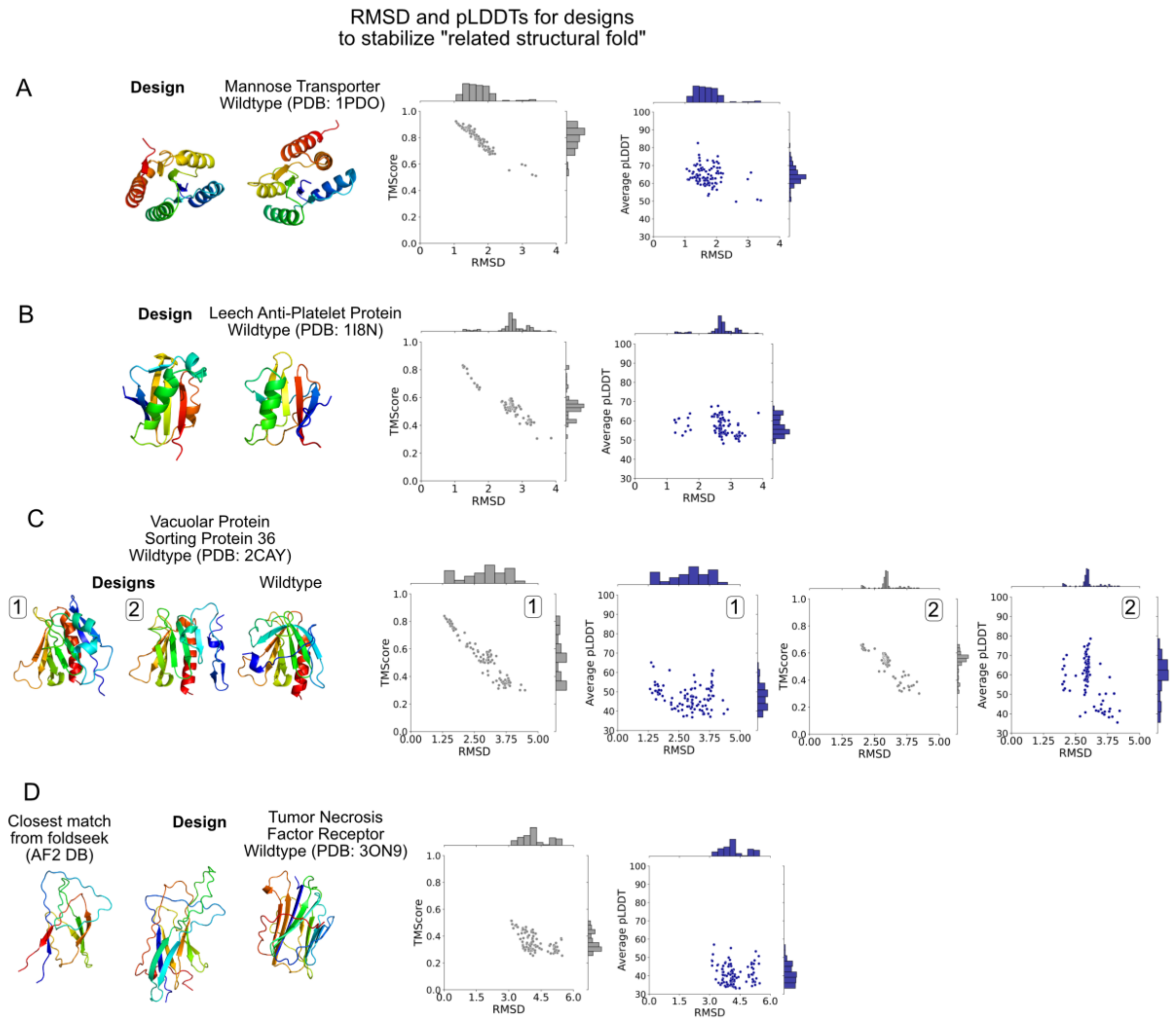


**SI Figure 6. Structural neighborhood of selected proteins from the TS50 test set. (A) 1PDO (B) 1I8N (C) 2CAY (D) 3ON9.**

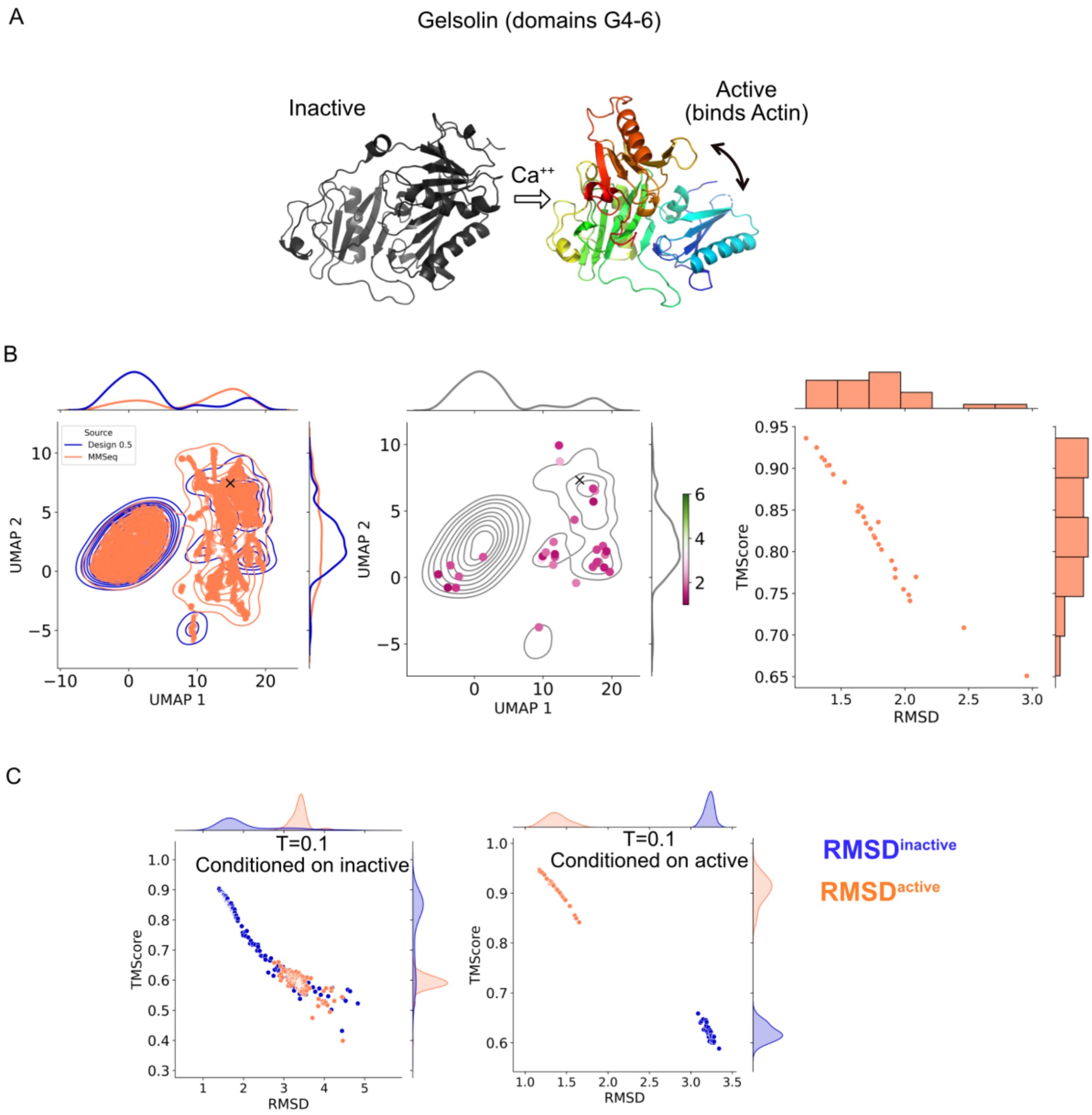


**SI Figure 7. (A**) Inactive and active conformations of Gelsolin. **(B)** (Left) Sampled sequences for the active state at T=0.5 (orange) with sequences from MMSeqs2 (blue). (Center) RMSD of selected sequences. (Right) RMSD of sampled sequences with respect to the active structure. **(C)** Distribution of RMSD and TMScore for designs conditioned on inactive and active conformations.

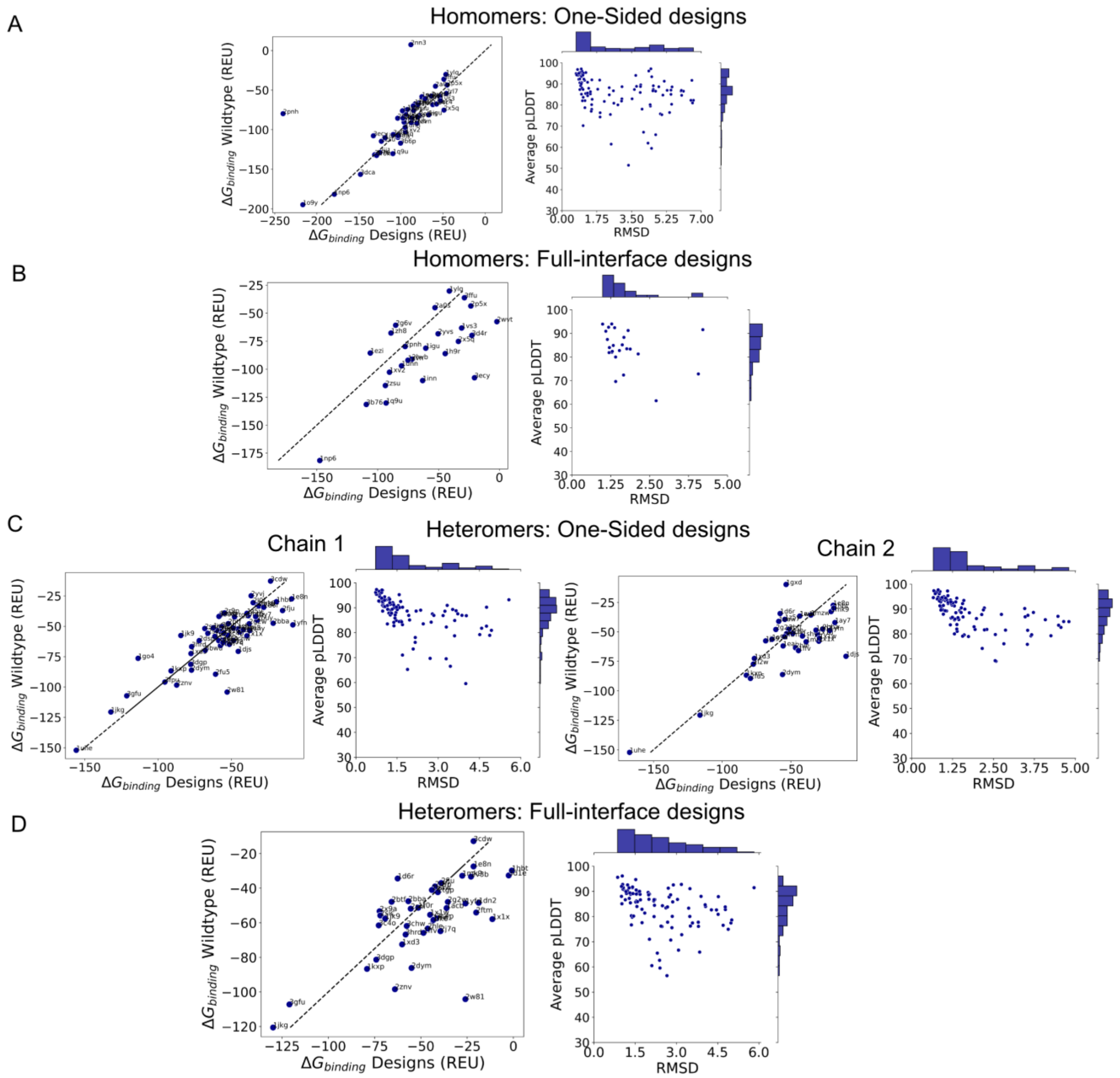


**SI Figure 8. RMSD, pLDDTs and comparison of binding energies for the PPI dataset. (A)** One-sided designs for the homomers. **(B)** Full interface designs for the homomers. **(C)** One-sided designs for the heteromers. **(D)** Full interface designs for the heteromers.

**SI Table 1. Set of 10 nanobodies in the AbAg70 dataset.**

| |
|---|
| 6ey6,4y7m,3rjq,6cwg,5hgg,5sv3,6dbg,4m3k,5jmo,2i25 |
